# Thermal proteome profiling reveals distinct target selectivity for differentially oxidized oxysterols

**DOI:** 10.1101/2021.01.05.425440

**Authors:** Cecilia Rossetti, Luca Laraia

## Abstract

Oxysterols are produced physiologically by many species, however their distinct roles in regulating human (patho)physiology have not been studied systematically. The role of differing oxidation states and sites in mediating their biological functions is also unclear. As individual oxysterols have been associated with atherosclerosis, neurodegeneration and cancer, a better understanding of their protein targets would be highly valuable. To address this, we profiled three A- and B-ring oxidized sterols as well as 25-hydroxycholesterol using thermal proteome profiling (TPP), validating selected targets with the cellular thermal shift assay (CETSA) and isothermal dose response fingerprinting (ITDRF). This revealed that the site of oxidation has a profound impact on target selectivity, with each oxysterol possessing an almost unique set of target proteins. However, overall targets clustered in pathways relating to vesicular transport and lipid metabolism and trafficking, suggesting that while individual oxysterols bind to a unique set of proteins, the processes they modulate are highly interconnected.

## Introduction

Dysregulation of cholesterol homeostasis is a severe condition leading to inadequate or excessive tissue cholesterol levels. Hypercholesterolemia has been identified as a common risk factor of diverse disorders, including breast, colorectal, prostatic and testicular cancer^[1]^ together with coronary, artery and Alzheimer's diseases.^[2],[3]^ Oxidative metabolites of cholesterol, termed oxysterols, contribute to the regulation of cholesterol homeostasis with different transcriptional and non-genomic mechanisms, which are still incompletely understood.^[4],[5],[6]^ Additionally, recent research suggests that they may play distinct roles not directly connected to the regulation of cholesterol homeostasis, including mediating membrane contact sites and trafficking. Evidence has also associated increased oxysterol levels to cancer progression, the mechanisms of which remain to be elucidated.^[7]^

Of the over twenty oxysterols identified, side-chain oxidized sterols and particularly 25-hydroxycholesterol (25-HC) have been the most widely studied. They have been shown to modulate the activity of cholesterol transport proteins and transcription factors involved in regulating cholesterol homeostasis. However, A- and B-ring oxidized sterols have been less well studied, in particular in relation to their target profile. Those oxidized at the C7 position, such as 7-ketocholesterol (7-KC), are most frequently detected at high levels in atherosclerotic plaques^[8]^ and in the plasma of patients with high cardiovascular risk factors.^[9]^ Furthermore, 7-KC displays toxicity at higher concentrations, accompanied by a pronounced effect on lysosomal activity.^[10]^ The precise mechanisms by which this occurs are still unknown. For oxysterols oxidized at 4-, 5- and 6-positions virtually no targets have been annotated, with the partial exception of the liver X receptor (LXR). Crucially, the effect on biological activity of different oxidative modifications on the sterol backbone has not been explored.

For all of the reasons above, the systematic discovery of oxysterol target proteins will be of profound importance in determining their (patho)physiological roles. Herein we describe the systematic identification of oxysterol target proteins using thermal proteome profiling (TPP). The oxidation site and state significantly affected the target profile for each oxysterol tested, with only two proteins identified as targets for more than one oxysterol. Of these, the vacuolar protein sorting associated protein 51 (VPS51) was validated more comprehensively as a protein that binds oxysterols. Though different, most oxysterol targets clustered in pathways and processes related to vesicular transport as well as lipid metabolism and transfer, and most targets were localized at intracellular membranes. These results suggest specific but different roles for individual oxysterols and provide a blueprint for further studies on these important metabolites.

## Results and Discussion

### Identification of oxysterol target proteins using thermal proteome profiling

To identify potential oxysterol target proteins, TPP was selected as the method of choice ^[11][12]^ (Figure 1A). This method is advantageous over other target identification methods as it does not require pre-functionalization, immobilization or modification of the compound of interest and has been shown to offer excellent proteome coverage. Furthermore, the use of selected detergents including NP-40, has successfully enabled the identification of a large proportion of membrane proteins, which is particularly relevant as this is where a large proportion of known sterol targets are located.^[13]^ We opted to carry out experiments in cell lysates for the primary screening efforts, for increased reproducibility^[14],[15]^ and data interpretation simplicity.^[16]^ The use of cell lysates enables the evaluation of direct oxysterol target engagement without additional sources of variability deriving from factors such as membrane transport, accumulation and cell metabolism, which are prominent in experiments with intact cells. We selected 4β-hydroxycholesterol (4β-HC), cholestane-3β,5α,6β-triol (CT) and 7-ketocholesterol (7-KC) as representative A/B-ring oxidized sterols which cover all four known oxidation sites and arise through enzymatic, but also spontaneous oxidation (Figure 1B). The oxysterols were also selected with the aim of elucidating how the different oxidation pattern on the sterol core determines the selectivity of these important metabolites. Furthermore, we also included 25-hydroxycholesterol (25-HC) as an oxysterol that has been more widely studied and applied, but whose complete target profile had also not been elucidated.

**Figure 1.**
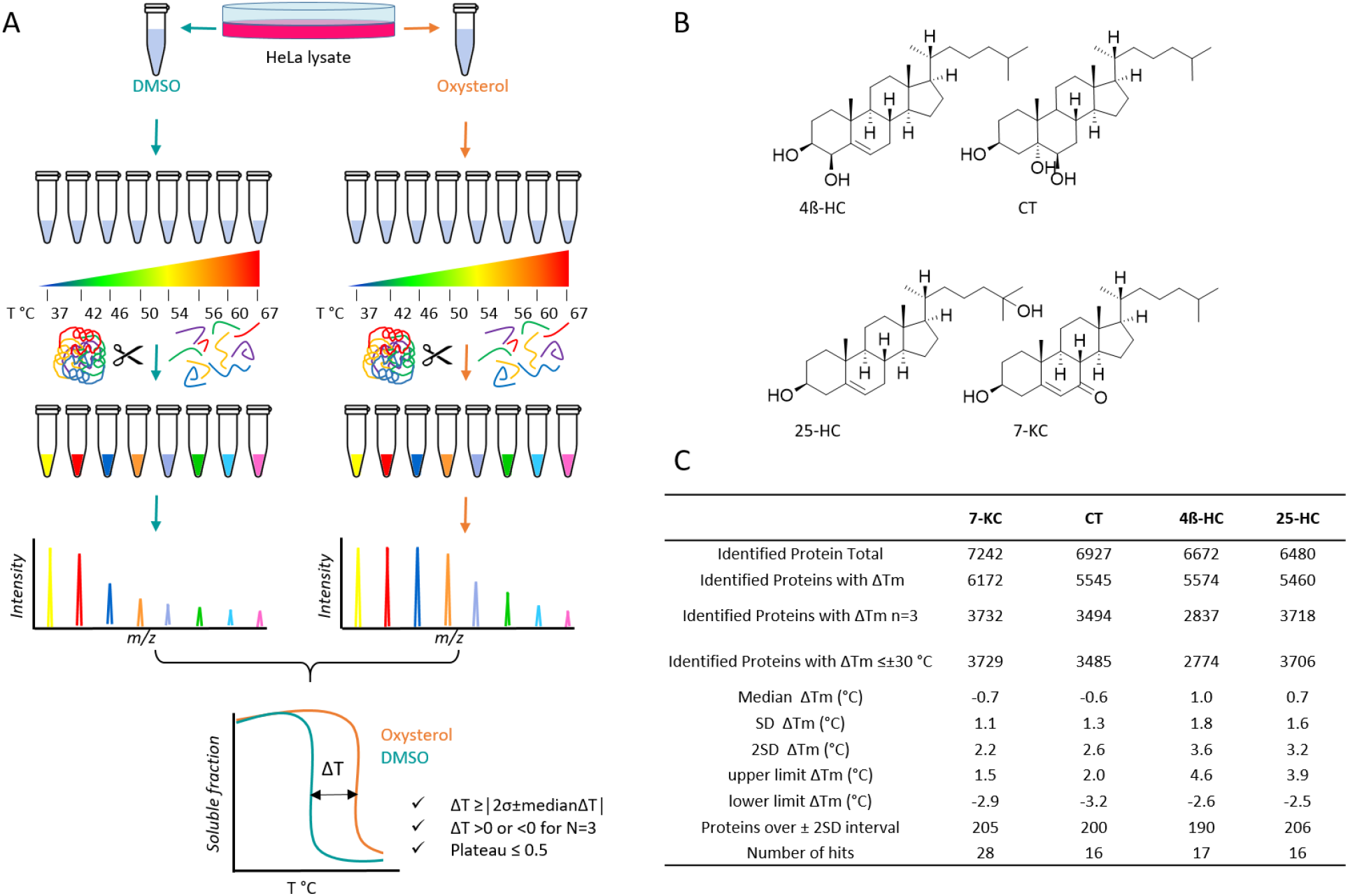
Target identification of oxysterols using thermal proteome profiling. A) Workflow of the thermal proteome profiling experiments and criteria for the identification of putative oxysterol targets; B) Structures of the tested oxysterols: 4ß-hydroxycholesterol (4ß-HC), cholestane-3β,5α,6β-triol (CT), 25-hydroxycholesterol (25-HC) and 7-ketocholesterol (7-KC); C) Summary table of the identified proteins from the HeLa proteome analysis and setting of the threshold limits for the identification of putative hits.

TPP enabled the identification and monitoring of changes in thermal stability of up to 7000 proteins, upon the incubation of HeLa cell lysates with the different oxysterols (Figure 1C). From these, it was possible to calculate thermal shifts for about 85% of the identified proteins. To define which shifts in melting temperatures were significant, two standard deviations from the median of all the calculated shifts was deemed appropriate, in line with previous reports.^[17]^ In the screening process, proteins with a significant change in melting temperature following oxysterol exposure were filtered according their melting curves normalized to the lowest temperature. Proteins displaying a shift in the same direction (positive or negative ΔT_m_) in all three replicates and with a curve plateau corresponding to a fraction of soluble protein less or equal to 0.5 were selected as potential targets (Figure 1A and 1C).

The entire screening set produced a list of 77 hits considered as putative targets for at least one of the tested oxysterols (Figure 2A). Overall, the re-identification of 10 known cholesterol binding proteins as determined by affinity-based probes^[18]^ (Supporting Information (SI) Table S1), both validates the use of TPP for identifying novel oxysterol target proteins, but also highlights the wealth of previously unidentified sterol interactors. Interestingly, the overlap of the candidate targets between the different oxysterols was remarkably low. Only 7-KC and 4β-HC shared two putative interacting proteins (Figure 2B). While this result may appear unexpected, it is in fact consistent with previously reported studies describing the binding of oxysterols to the cholesterol transport proteins Niemann Pick Class 1 (NPC1) and Aster-A (also known as GRAMD1A). Both were shown to bind 25-HC, but displayed no binding of A- or B-ring oxidized sterols.^[19][20][21]^ These examples and our data suggest that these particular oxysterols do not exert their function by modulating traditional (oxy)sterol-associated proteins. Crucially, our data suggests that different oxysterols show distinct target profiles that are dependent on the position and level of oxidation.

**Figure 2.**
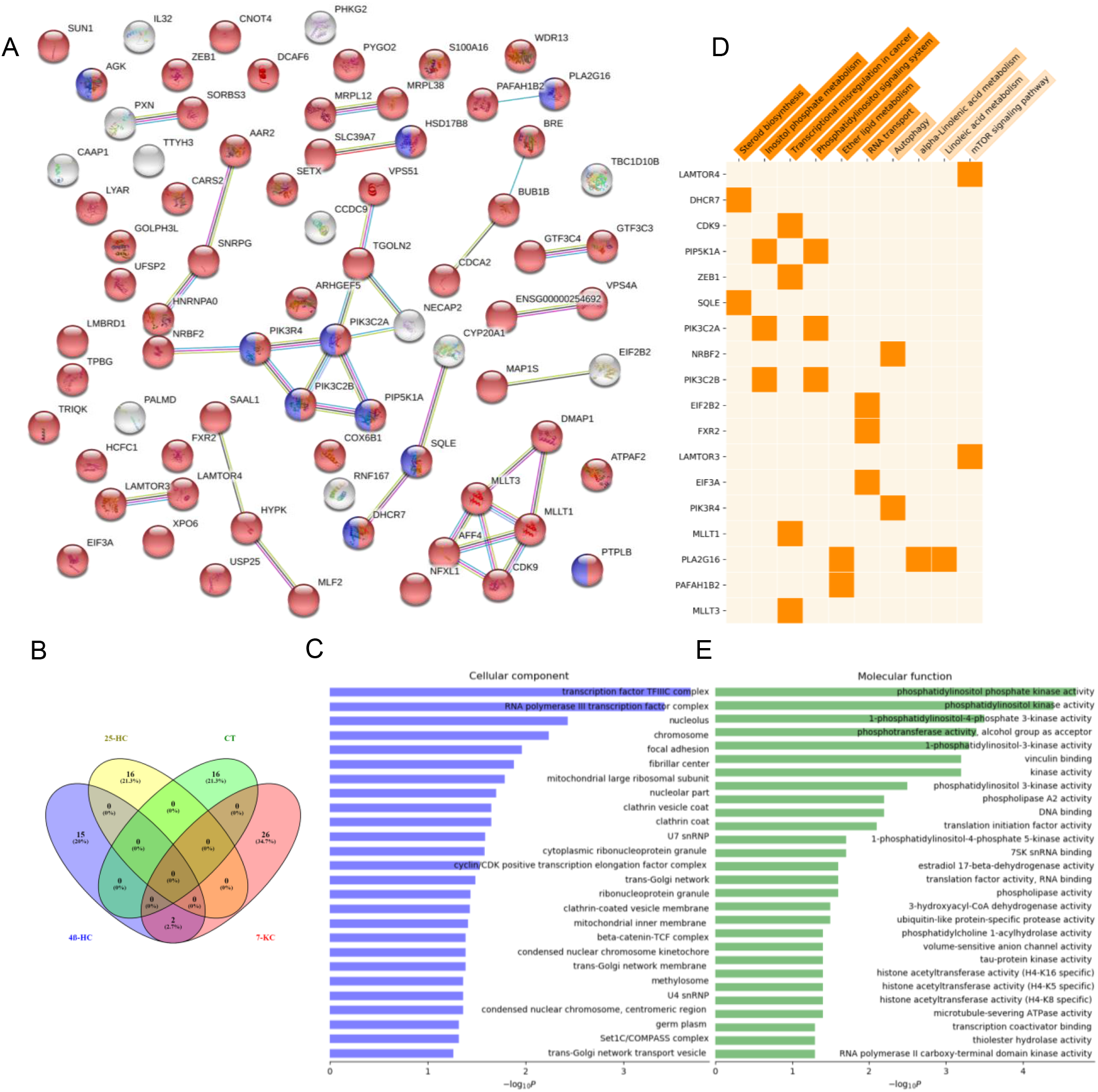
Analysis of all the putative targets for the tested oxysterols 4ß-HC, 25-HC, CT and 7-KC. A) STRING functional analysis with proteins from intracellular membrane-bounded organelle highlighted in red (GO:0043231; FDR: 5.20e-06), and proteins involved in the metabolism of lipids in blue (HSA-556833; FDR: 0.027); B) Venn diagram of the putative targets for each of the oxysterol and overlap among common targets; C) GO Cellular Components enriched from the analysis of all the putative targets; D) KEGG Pathway analysis and target contribution for each pathway. Pathways are colored according their significance from orange to white to indicate p-values from 0.002 to 0.1; E) GO Molecular Functions enriched from the analysis of all the putative targets.

Despite marked differences in their individual target profiles, some general trends could be observed clearly. The functional enrichment analysis of the identified candidate targets (Figure 2A) showed an enrichment of the intracellular membrane compartments (highlighted in red and listed in SI Table S2). Of these, several proteins associated with clathrin coated vesicle (CCV) transport were identified (Figure 2C). CCV transport is known to require cholesterol,^[22]^ however except for OSBP the mediators of this effect were unknown, and oxysterols were not suggested to modulate this process. Trans-Golgi network (TGN) membrane associated proteins were also significantly targeted. As CCVs are known to also form at the TGN, this could suggest an overall link between oxysterols and CCV transport. Unexpectedly, a large proportion of the RNA polymerase III transcription complex was identified as putative oxysterol targets (Figure 2C). In particular, constituents of the super elongation complex (SEC) were highly enriched, with 4β-HC targeting several of the components (*vide infra*).

Perhaps unsurprisingly, among the Reactome pathways enriched in the STRING analysis, the metabolism of lipids was identified (Figure 2A, highlighted in blue, and SI Table S3). This was due to the presence of known sterol biosynthetic and metabolic proteins but also by a large number of lipid kinases of different classes. In particular phosphatidyl inositol kinases (PIKs), which were targeted by multiple oxysterols, contributed to the enrichment of Phosphatidylinositol metabolism in both Reactome and KEGG pathways (SI table S3 and Figure 2C, respectively) and contributed to the enrichment of the Phosphatidylinositol (phosphate) kinase activity as the most significant molecular function in the GO analysis (Figure 2E). Proteins that regulated the mechanistic target of rapamycin complex 1 (mTORC1) either directly or indirectly were also abundant, confirming its essential role in regulating lipid metabolism.

### Proteome-wide profiling of 7-KC

We focused our initial analysis of specific oxysterol target proteins with 7-KC, as it is the most prominent and toxic of the non-enzymatically produced oxysterols. Several known and novel 7-KC targets with significant ΔT_m_ (Figure 3A and 3B) were identified. For example, squalene monooxygenase (SQLE) is a key cholesterol biosynthetic enzyme.^[5]^ 7-KC has been previously shown to lead to SQLE degradation, analogously to cholesterol and other sterols oxidized at the 7-position. The identification of a known 7-KC target protein further confirms that TPP is a suitable approach for oxysterol target protein identification. Importantly, destabilized proteins were considered as putative targets (SI Table S4). Among them, BRISC and BRCA1-A complex member 2 (BRE), is known to be involved in the defective synthesis of steroid hormones and accumulation large quantities of cholesterol under stress or under the influence of steroid hormones.^[23],[24]^ Among other stabilized proteins, several are involved in PI metabolism, including PIP5K1A, which is the main source of cellular PI4,5P_2_. Nuclear receptor-binding factor 2 (NRBF2) is known to modulate PI3K-III activity by stabilization of the VPS34 complex I, a key autophagy-related kinase.^[25]^

**Figure 3.**
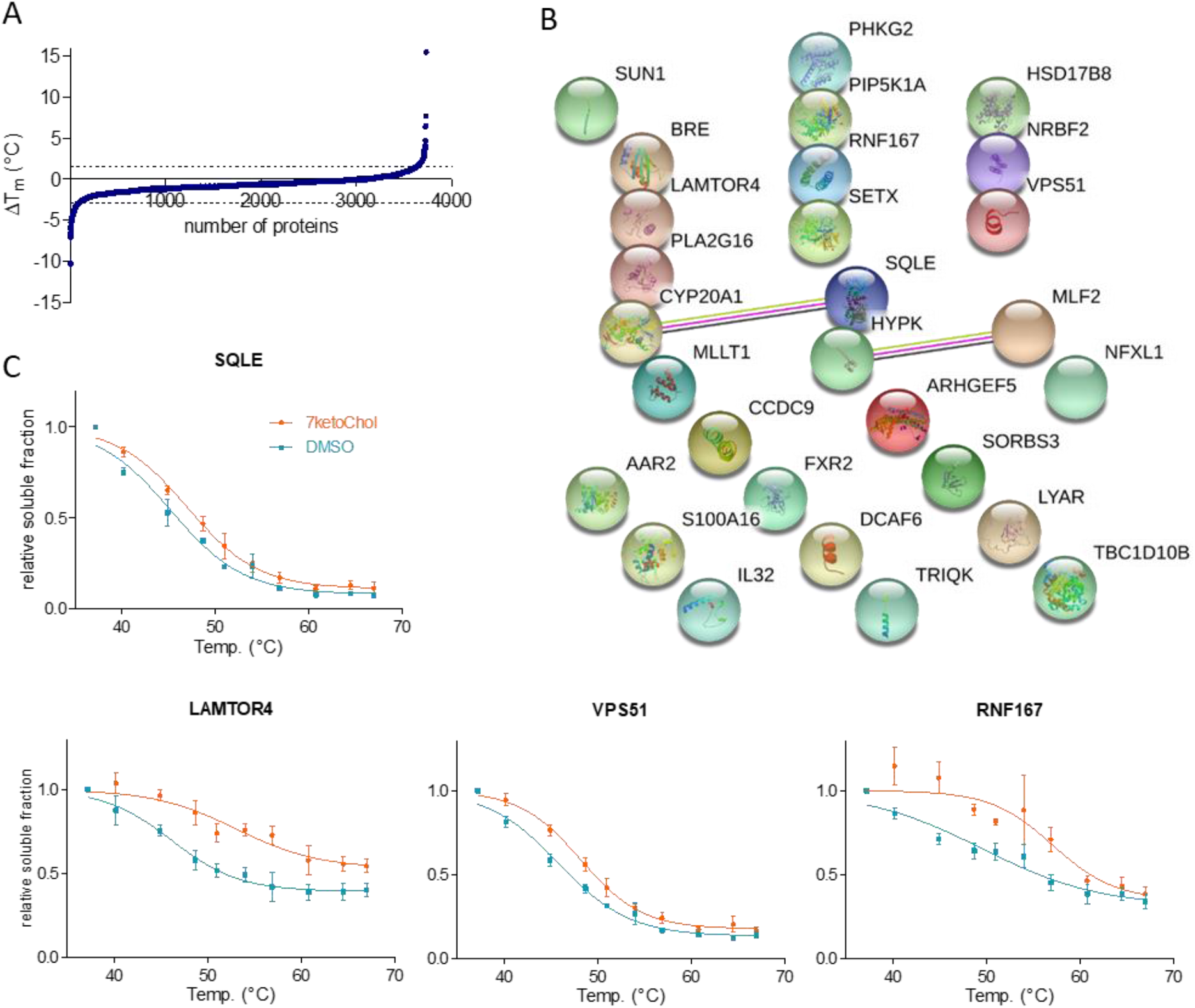
TPP analysis of 7-KC. A) Melting temperature shifts of the entire HeLa proteome. Significant shifts lies outside the 2 standard deviation interval marked with dotted lines. B) STRING functional analysis of the putative targets selected from the TPP screening assay. C) Melting curves of Squalene monooxygenase (SQLE), E3 ubiquitin-protein ligase RNF167 (RNF 167), Vacuolar protein sorting-associated protein 51 homolog (VPS51) and the Ragulator protein complex protein LAMTOR4. Data is mean ± sem of three independent experiments.

The two targets most stabilized by 7-KC were associated with lysosomal functions (SI Table S4). E3 ubiquitin-protein ligase RNF167 and Ragulator complex protein LAMTOR4 are both localized in lysosomes, where they perform ubiquitin protein ligase activity and regulation of TOR signaling activity, respectively (see Figure 3C for associated melting curves). RNF167 has been found to regulate AMPA receptor-mediated synaptic transmission^[26]^ while the LAMTOR complex regulates mTOR signaling and thus cellular lipid metabolism more generally. The modulation of these targets may begin to explain the phenotypic effects elicited by 7-KC^[10]^. Accumulation of 7-KC in the lysosomes is thought to alter pH maintenance, reducing their ability to hydrolyze and process cellular debris, as it has already described for lysosomal accumulation of cholesterol^[27]^.

The presence of VPS51 among the most stabilized proteins was intriguing, as it was one of only two (the other being AAR2 splicing factor homolog) proteins identified as putative new targets for more than one oxysterol (7-KC and 4β-HC). VPS51 is part of the Golgi-associated retrograde protein (GARP) complex, which is known to regulate cholesterol transport between early and late endosomes and the trans-Golgi network (TGN) via lysosomal NPC2^[28]^. However, a direct interaction of VPS51 with cholesterol, or indeed any sterol, had not been reported. Thus we selected VPS51 for further validation. The TPP results were initially validated by means of a cellular thermal shift assay (CETSA), with Western blot read-out. For both 7-KC and 4β-HC, we were able to reproduce the stabilization observed in the TPP experiment (Figure 4A and 4B, respectively), although the thermal shift was less pronounced for 4β-OHC. To address this discrepancy, we carried out an isothermal dose-response fingerprinting (ITDRF) experiment, which showed that 4β-HC stabilized VPS51 in a dose-dependent manner at 51 °C, confirming their putative interaction (Figure 4C).

**Figure 4.**
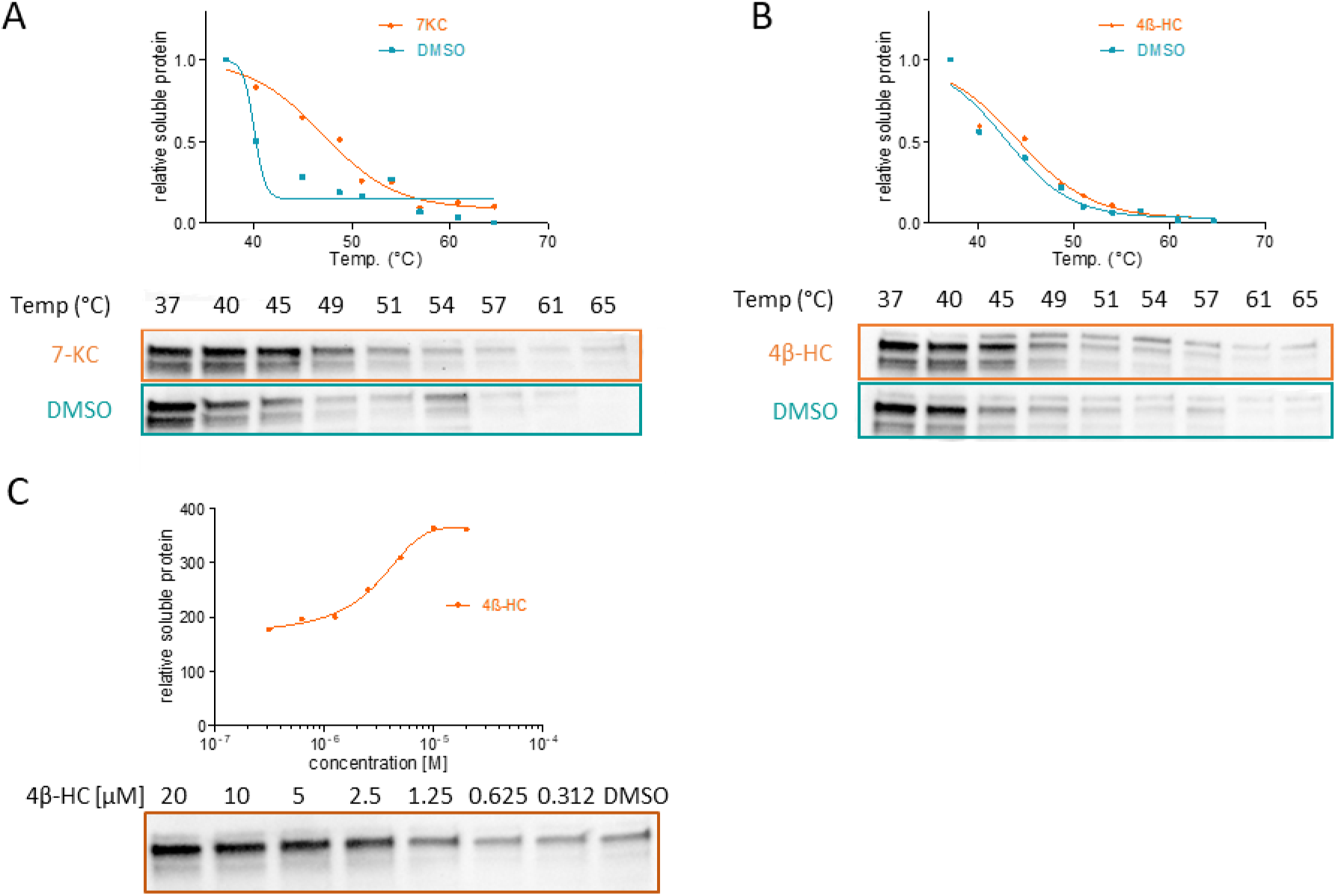
Target validation of VPS51. A) CETSA experiments for the validation of VPS51 with 7-KC; B) CETSA experiments for the validation of VPS51 with 4ß-HC; C) ITDRF experiment for the validation of VPS51 in 4ß-HC with related dose-response curve. Both reported isoforms of VPS51 are visible. Data is the mean of two independent experiments, representative blots are shown.

### Putative targets of 4β-hydroxy cholesterol

In addition to VPS51, 4β-HC appeared to target cholesterol transport in other ways (SI Table S5 and Figure S2). Targets were clearly enriched vacuolar transport processes, including VPS51, VPS4A and PIK3R4 (VPS15) (Figure S2, green). VPS15 is a component of the class III PI3K complex, which is a key component of autophagy initiation, strengthening the link observed with other oxysterols. VPS4A has been extensively associated with cholesterol transport, in a function not directly governed by its role in disassembling the endosomal sorting complex required for transport (ESCRT-III) polymer.^[29][30]^

The stabilization of translation initiation factors EIF3A and EIF2B, may be connected to the more general targeting of other mTOR regulators including LAMTOR3 and 4 by oxysterols, since it has been shown that the mTOR complex mediates assembly of the translation preinitiation complex (PIC) modulating the function of EIF3 in the translation of mRNAs encoding proteins.^[31]^ Very recently 4β-HC has been shown to act as a pro-lipogenic factor by enhancing Sterol Regulatory Element Binding Protein 1c (SREBP1c) expression in an LXR-dependent manner.^[32]^ In this context, we found that 4β-HC (de)stabilized a series of transcriptional regulators, including the general transcription factor 3C (GTF3C) and two components of the super elongation complex, cyclin-dependent kinase 9 (CDK9) and AF4/FMR2 family member 4 (AFF4). This raises the possibility that transcriptional elongation of SREBP1c may require 4β-HC’s ability to interact with the SEC.

### Putative targets of 25-hydroxy cholesterol

Putative targets of 25-HC were strongly enriched in PI metabolism (SI Figure S3). Stabilization of the Phosphatidylinositol 4-phosphate 3-kinase C2 domain-containing subunit alpha (PIK3C2A) and destabilization of related subunit beta (PIK3C2B), allowed the identification of two of the three isoforms of the class II PI3Ks. These known to play key roles in clathrin-mediated endocytosis.^[33]^ The ability of 25-HC to modulate PIK3C2A was tested in an enzymatic kinase profiling assay; however no change in kinase activity was observed (SI Table S8). This does not necessarily de-validate the target, as binding in an allosteric pocket may modulate protein-protein interactions rather than enzymatic activity. Similarly, all other putative kinase targets of oxysterols (CDK9, PHKG2 and PIP5K1A), were tested with the related assays without showing significant increase or decrease in enzymatic activity (SI Table S9-S11).

Unsurprisingly, 25-HC also stabilized regulators of cholesterol biosynthesis and metabolism, including 7-dehydrocholesterol reductase (DHCR7). DHCR7 catalyzes the last step in the biosynthesis of cholesterol and when mutated has been associated with the developmental disease Smith-Lemli-Opitz syndrome.^[34]^ Binding to oxysterols such as 7-KC is known to induce its proteasomal degradation; however, interestingly this effect was not reported for 25-HC.^[35]^ The stabilization of Host cell factor 1 (HCFC1) by 25-HC could link the regulation of an intragenic region in the HCFC1 gene by Sterol regulatory element-binding protein 1 (SREBP1),^[36]^ which is itself regulated by cholesterol and 25-HC.^[37]^ Finally, 25-HC targeted the Ragulator complex protein LAMTOR3, and the Transmembrane 9 superfamily member 1 (ENSG00000254692), a protein involved in authophagy,^[38]^ which is known to be regulated by cholesterol metabolism.^[39]^

### Putative targets of cholestane triol

CT targets were significantly enriched in autophagic and golgi-associated proteins (SI Figure S4). The most stabilized protein was Paxillin (PXN), an autophagy substrate that interacts with LC3 during focal adhesion (FAs) disassembly in highly metastatic tumor cells.^[40]^ FA turnover is reportedly influenced by ORP3-mediated lipid exchange,^[41]^ which may explain the association of oxysterols with PXN. A significantly destabilized target was the microtubule-associated protein 1S (MAP1S), whose deficiency causes impaired autophagic degradation of lipid droplets, which then accumulate in normal renal epithelial cells, initiating the development of renal cell carcinomas.^[42]^

Golgi phosphoprotein 3-like (GOLPH3L) was the most destabilized putative target. Interestingly, this protein is also associated to the AKT/mTOR pathway, since it contributes to the tumorigenesis of Hepatocellular carcinoma increasing cell proliferation by the activation of mTOR signaling via overexpression of mTORC1.^[43]^ Adaptin ear-binding coat-associated protein 2 (NECAP2) promotes fast endocytic recycling of epidermal growth factor receptor (EGFR) and of the tumor necrosis factor receptor (TfnR) through the recruitment of AP-1–clathrin machinery to early endosomes. In order to facilitate the receptor recycling, early endosomes receive endocytosed material from clathrin-dependent and -independent pathways and sort cargo for recycling to the cell surface, retrograde transport to the Golgi or degradation in lysosomes.^[44]^ NECAP2 sits at a node in the overall oxysterol target interaction map, and would thus be an intriguing target for further study.

## Conclusion

In summary, we have carried out the first systematic exploration of oxysterol target proteins using thermal protein profiling as the enabling technology. TPP proved convenient for screening small compounds sets such as the four oxysterols we selected, as it does not require compound modification or functionalization. Furthermore, previously identified sterol-binding proteins were re-identified here, validating the approach.^[18]^ Strikingly, our results demonstrate that oxysterols which differ from cholesterol by the addition of just one or two oxygen atoms, display distinct target profiles, with only two proteins identified as targets of more than one oxysterol. To the best of our knowledge this has never been conclusively shown or systematically studied.

Although virtually no overlap between the oxysterol targets was present, targets were enriched in lipid metabolism, mTOR signaling, vesicle trafficking and transcriptional regulators. The intracellular membrane localization of most target proteins is also consistent with the lipophilic nature of the compounds, and their reported membrane association. Of the two proteins which share two oxysterols as putative targets, VPS51 was further validated using CETSA and ITDRF experiments. Although its role in mediating cholesterol transport by targeting NPC2 to the lysosomes as part of the GARP complex is known, our data raises the intriguing possibility that this event is regulated by (oxy)sterols themselves. The specific target profiles of the individual oxysterols studied may also begin to explain the phenotypes they induce. In particular 7-KC has previously been shown to affect lysosomal integrity and activity. The fact that several of the putative targets identified are lysosomal membrane proteins may begin to offer an explanation for this observed effect. Importantly, future work to determine whether target (de)stabilization by oxysterols occurs through direct binding or is mediated by a complex will be necessary.

To conclude, TPP is a robust technology to identify new oxysterol target proteins, and the data provided herein provides an extensive resource as well as a wealth of testable hypotheses linking oxysterols to lipid metabolism and transport, vesicle trafficking and transcription. Despite the exciting developments achievable with this technique, it is important to note that like all target identification methods, TPP also has its caveats. False negatives are more common with this technique as target proteins may not be (de)stabilized by small molecules they interact with, or that very high compound concentrations are required to see observe a meaningful effect. This was particularly apparent for known 25-HC protein targets including OSPB, NPC1 and certain STARDs which were not identified as putative targets although they are present in our MS data and more generally in meltome analyses. ^[45]^ OSBP, STARD2, NPC1 and NPC2 proteins were identified in the HeLa cell proteome, but their thermal shift was not considered significant according the chosen criteria or was not determined in all three replicates. In this regard, the arbitrary exclusion of proteins with shifts lower than two standard deviations from the median might particularly affect the recognition of protein targets belonging to compounds whose meltome more generally altered from the DMSO control. While the use of NP-40 facilitates recovery of membrane proteins, it has recently been shown that different detergent types and concentrations can affect which proteins are recovered in the final analysis, introducing a slight bias.^[13]^ Despite this, we believe that TPP and its variants including ITDRF will be applied increasingly for (off)-target identification and validation.

## Supporting information

Supplementary figures and experimental procedures

## Acknowledgements

We would like to thank Assoc. Prof. Erwin Schoof from DTU Proteomics Core for excellent advice and support and Prof. Ulrich auf dem Keller for access to cell culture and reagents at DTU Bioengineering. We would also like to thank Dr. Petra Janning and Malte Metz for invaluable advice regarding the data analysis. We would also like to acknowledge the Novo Nordisk Foundation (NNF17OC0028366) and DTU for funding.

