## Supplementary figures and experimental procedures for "Thermal proteome profiling reveals distinct target selectivity for differentially oxidized oxysterols"

#### Supplementary Material

### Supporting Information

*Table S1 List of candidate targets from TPP also identified in the sterol-pull down experiment described in Hulce et al.*[1]

| *Gene Id* | *Protein Name* |
|---|---|
| *PTPLB* | Very-long-chain (3R)-3-hydroxyacyl-CoA dehydratase 2 |
| *DHCR7* | 7-dehydrocholesterol reductase |
| *TTYH3* | Protein tweety homolog 3 |
| *SLC39A7* | Zinc transporter SLC39A7 |
| *AGK* | Isoform 1 of Acylglycerol kinase, mitochondrial |
| *TPBG* | Trophoblast glycoprotein precursor |
| *NFXL1* | Isoform 1 of NF-X1-type zinc finger protein |
| *CYP20A1* | Cytochrome P450 family 20subfamily A member 1 |
| *SUN1* | SUN domain-containing protein 1 |
| *SQLE* | Squalene monooxygenase |

*Table S2 Cellular Component enriched in the STRING Protein-Protein Interaction Networks Functional Enrichment Analysis analysis for all the putative targets found*

| *Cellular Component* | *Description* | *FDR* |
|---|---|---|
| *GO:0043231* | intracellular membrane-bounded organelle | 5.20E-06 |
| *GO:0043227* | membrane-bounded organelle | 3.37E-05 |
| *GO:0005622* | intracellular | 0.00013 |
| *GO:0005634* | nucleus | 0.00013 |
| *GO:0005737* | cytoplasm | 0.00013 |
| *GO:0005942* | phosphatidylinositol 3-kinase complex | 0.00013 |
| *GO:0031981* | nuclear lumen | 0.00013 |
| *GO:0043229* | intracellular organelle | 0.00013 |
| *GO:0070013* | intracellular organelle lumen | 0.00013 |
| *GO:1990234* | transferase complex | 0.00052 |
| *GO:0032991* | protein-containing complex | 0.00057 |
| *GO:0005829* | cytosol | 0.0011 |
| *GO:0008023* | transcription elongation factor complex | 0.0012 |
| *GO:0071986* | Ragulator complex | 0.0054 |
| *GO:0061695* | transferase complex, transferring phosphorus-containing groups | 0.006 |
| *GO:0000127* | transcription factor TFIIIC complex | 0.0065 |
| *GO:0035032* | phosphatidylinositol 3-kinase complex, class III | 0.0065 |
| *GO:0005623* | cell | 0.0102 |
| *GO:0005654* | nucleoplasm | 0.0109 |

| | | |
|---|---|---|
| *GO:0005730* | nucleolus | 0.0109 |
| *GO:0043232* | intracellular non-membrane-bounded organelle | 0.0171 |

*Table S3 Reactome Pathways enriched in the STRING Protein-Protein Interaction Networks Functional Enrichment Analysis analysis for all the putative targets found*

| *Reactome Pathways* | *Description* | *FDR* |
|---|---|---|
| *HSA-112382* | Formation of RNA Pol II elongation complex | 2.70E-02 |
| *HSA-1483255* | PI Metabolism | 2.70E-02 |
| *HSA-1483257* | Phospholipid metabolism | 0.027 |
| *HSA-556833* | Metabolism of lipids | 0.027 |
| *HSA-674695* | RNA Polymerase II Pre-transcription Events | 0.027 |
| *HSA-1660517* | Synthesis of PIPs at the late endosome membrane | 0.0379 |
| *HSA-1660499* | Synthesis of PIPs at the plasma membrane | 0.0389 |

*Table S4 List of candidate targets from TPP analysis of 7keto-cholesterol (7-KC) with related thermal shifts ($\Delta T_m$). Values are the mean of three independent experiments.*

| *Gene Id* | *Protein Name* | *$\Delta T_m$ (°C)* |
|---|---|---|
| *RNF167* | E3 ubiquitin-protein ligase RNF167 | 7.7 |
| *LAMTOR4* | Ragulator complex protein | 6.5 |
| *MLLT1* | Protein ENL | 3.7 |
| *IL32* | Interleukin-32 | 2.9 |
| *VPS51* | Vacuolar protein sorting-associated protein 51 homolog | 2.8 |
| *TRIQK* | Triple QxxK/R motif-containing protein | 2.6 |
| *SUN1* | SUN domain-containing protein 1 (Fragment) | 2.5 |
| *DCAF6* | DDB1- and CUL4-associated factor 6 | 2.4 |
| *NRBF2* | Nuclear receptor-binding factor 2 | 2.2 |
| *TBC1D10B* | TBC1 domain family member 10B | 2.1 |
| *SQLE* | Squalene monooxygenase | 2.0 |
| *LYAR* | Cell growth-regulating nucleolar protein | -2.9 |
| *S100A16* | Protein S100-A16 | -3.0 |
| *AAR2* | Protein AAR2 homolog | -3.0 |
| *SORBS3* | Vinexin | -3.1 |
| *ARHGEF5* | Rho guanine nucleotide exchange factor 5 | -3.1 |
| *BRE* | BRISC and BRCA1-A complex member 2 | -3.2 |
| *FXR2* | Fragile X mental retardation syndrome-related protein 2 | -3.4 |
| *HSD17B8* | Estradiol 17-beta-dehydrogenase 8 | -3.7 |
| *NFXL1* | NF-X1-type zinc finger protein | -3.8 |
| *SETX* | Probable helicase senataxin | -3.9 |
| *CYP20A1* | Cytochrome P450 family 20subfamily A member 1 | -4.0 |
| *MLF2* | Myeloid leukemia factor 2 | -4.0 |
| *HYPK* | Huntingtin-interacting protein | -4.2 |
| *PIP5K1A* | Phosphatidylinositol 4-phosphate 5-kinase type-1 alpha | -4.7 |

| | | |
|---|---|---|
| *PLA2G16* | HRAS-like suppressor 3 | -4.7 |
| *PHKG2* | Phosphorylase b kinase gamma catalytic chain, liver/testis isoform | -4.7 |
| *CCDC9* | Coiled-coil domain-containing protein 9 | -5.0 |

*Table S5 List of candidate targets from TPP analysis of 4ß-hydroxycholesterol (4ß-HC) with related thermal shifts ($\Delta T_m$). Values are the mean of three independent experiments.*

| *Protein gene id* | *Name* | *$\Delta T_m$ (°C)* |
|---|---|---|
| *CDK9* | Cyclin-dependent kinase 9 | 6.4 |
| *VPS4A* | Vacuolar protein sorting-associated protein 4A | 6.4 |
| *VPS51* | Vacuolar protein sorting-associated protein 51 | 6.1 |
| *GTF3C4* | General transcription factor 3C polypeptide 4 | 6.1 |
| *USP25* | Ubiquitin carboxyl-terminal hydrolase 25 | 5.7 |
| *EIF3A* | Eukaryotic translation initiation factor 3 subunit A | 5.6 |
| *AFF4* | AF4/FMR2 family member 4 | 5.5 |
| *XPO6* | Exportin-6 | 5.3 |
| *EIF2B2* | Translation initiation factor eIF-2B subunit beta | 5.1 |
| *PIK3R4* | Phosphoinositide 3-kinase regulatory subunit 4 | 4.9 |
| *AGK* | Acylglycerol kinase, mitochondrial | 4.9 |
| *PTPLB* | Very-long-chain (3R)-3-hydroxyacyl-CoA dehydratase 2 | 4.7 |
| *SAAL1* | Protein SAAL1 | 4.6 |
| *MRPL38* | 39S ribosomal protein L38, mitochondrial | -2.7 |
| *AAR2* | Protein AAR2 homolog | -2.8 |
| *GTF3C3* | General transcription factor 3C polypeptide 3 | -4.0 |
| *PAFAH1B2* | Platelet-activating factor acetylhydrolase IB subunit beta | -7.5 |

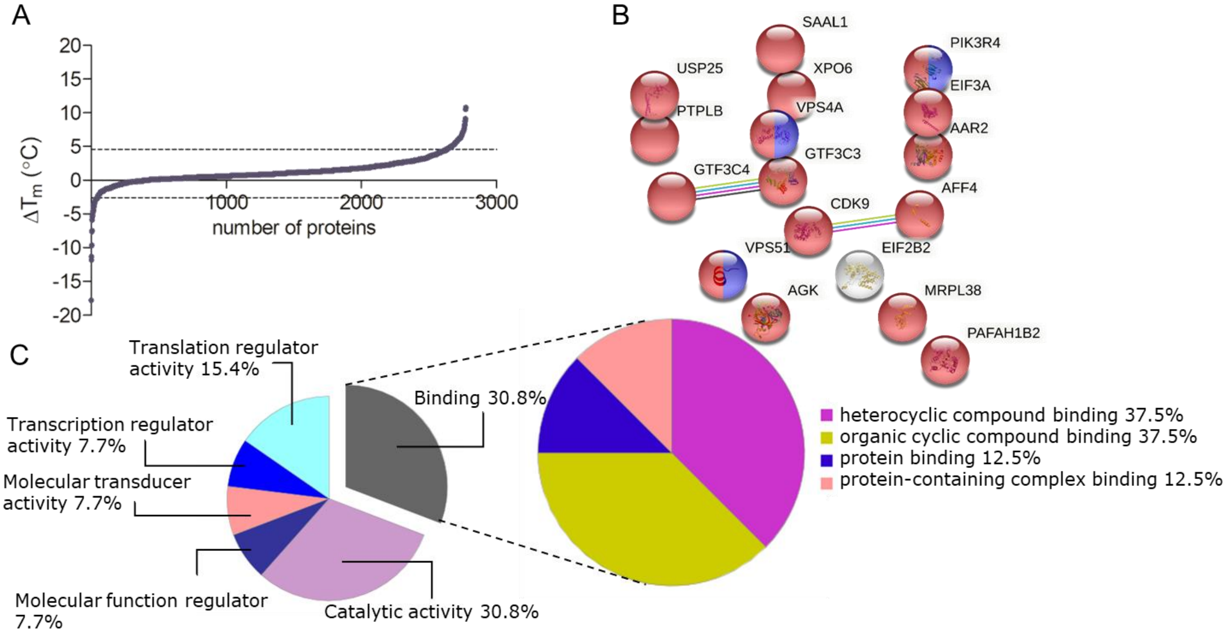

*Figure S1 A) Distribution of the melting temperature shifts for 4ß-HC and two standard deviation range (dashed line); B) STRING functional enrichment analysis of the candidate targets of 4ß-HC. Among the biological process here are shown the amide biosynthetic process (GO:0043604 in blue) and the vacuolar transport (GO: 0007034 in green) enriched with FDR values of 0.0143 and 0.0144 respectively. In red are the putative targets belonging to the cellular component intracellular membrane-bounded organelle (GO:0043231) enriched with a FDR of 0.0149; C) Pie chart displaying the molecular function of the candidate targets of 4ß-HC obtained by Panther classification system. On the left, is reported the zoom of the biding activity percentage (detached from the smaller pie chart) with related legend and percentage for each function contributing to the binding GO term.*

*Table S6 List of candidate targets from TPP analysis of 25-hydroxycholesterol (25-HC) with related thermal shifts ($\Delta T_m$). Values are the mean of three independent experiments.*

| *Gene Id* | *Protein Name* | $\Delta T_m$ (°C) |
|---|---|---|
| *HCFC1* | Host cell factor 1 | 9.7 |
| *ENSG00000254692* | Transmembrane 9 superfamily member 1 | 6.2 |
| *PYGO2* | Pygopus homolog 2, isoform CRA_b | 5.7 |
| *WDR13* | WD repeat-containing protein 13 | 4.9 |
| *SNRPG* | Small nuclear ribonucleoprotein G | 4.9 |
| *MLLT3* | Protein AF-9 | 4.8 |
| *PALMD* | Palmdelphin | 4.7 |
| *CDCA2* | Cell division cycle-associated protein 2 | 4.2 |
| *COX6B1* | Cytochrome c oxidase subunit 6B1 | 4.1 |
| *TTYH3* | Protein tweety homolog 3 | 4.1 |
| *PIK3C2A* | Phosphatidylinositol 4-phosphate 3-kinase C2 domain-containing subunit alpha | 4.1 |
| *CAAP1* | Caspase activity and apoptosis inhibitor 1 | 4.1 |
| *CNOT4* | CCR4-NOT transcription complex subunit 4 | 4.1 |
| *LAMTOR3* | Ragulator complex protein LAMTOR3 | 4.0 |

| | | |
|---|---|---|
| *PIK3C2B* | Phosphatidylinositol 4-phosphate 3-kinase C2 domain-containing subunit beta | -2.5 |
| *DHCR7* | 7-Dehydrocholesterol reductase | -5.3 |

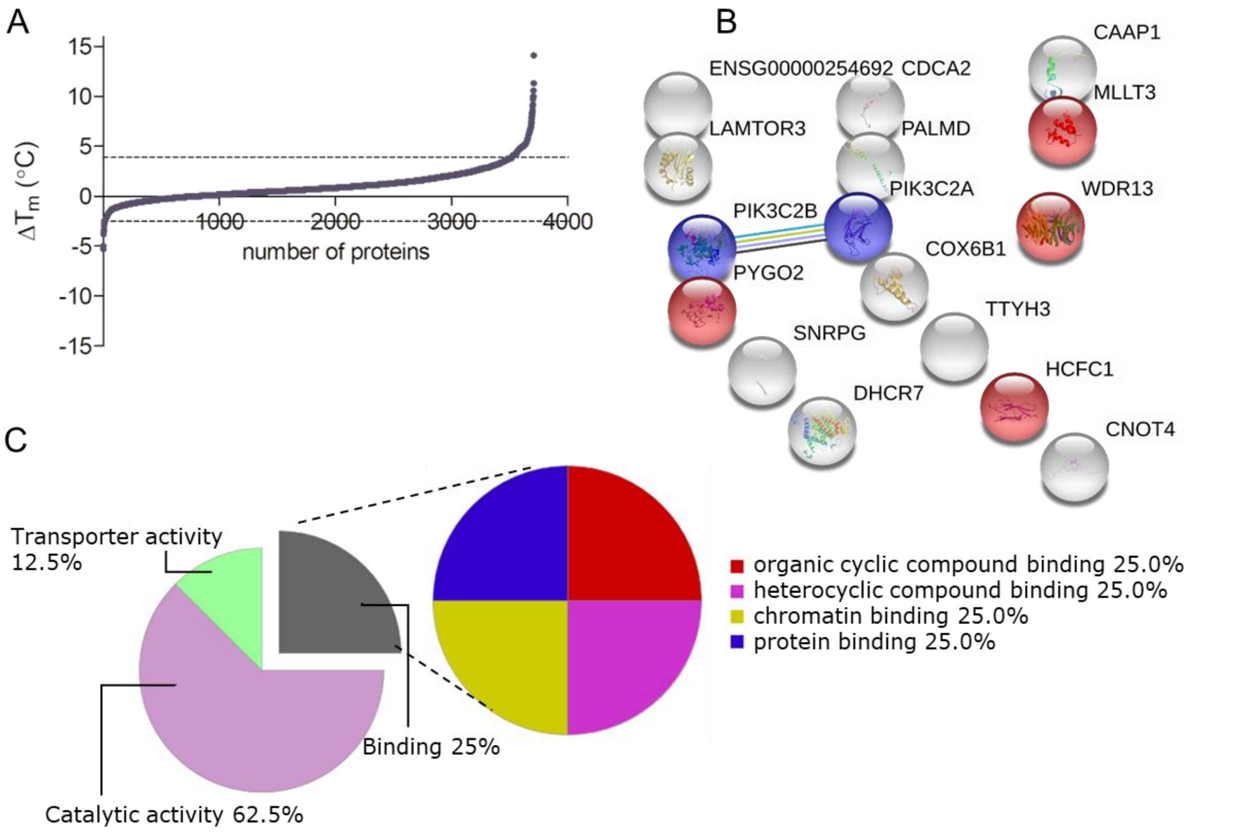

*Figure S2 A)Distribution of the melting temperature shifts for 25-HC and two standard deviation range (dashed line); B) STRING functional enrichment analysis of the candidate targets of 25-HC. The enriched molecular functions were the 1-phosphatidylinositol-4-phosphate 3-kinase activity (GO: 0035005 in blue) and the chromatin binding (GO:0003682 in red) with their respective FDRs of 0.0014 and 0.0101; C) Pie chart displaying the molecular function of the candidate targets of 25-HC obtained by Panther classification system. On the left, is reported the zoom of the biding activity percentage (detached from the smaller pie chart) with related legend and percentage for each function contributing to the binding GO term.*

*Table S7 List of candidate targets from TPP analysis of cholestane-3β,5α,6β-triol (CT) with related thermal shifts ($\Delta T_m$). Values are the mean of three independent experiments.*

| *Gene Id* | *Protein Name* | *$\Delta T_m$ (°C)* |
|---|---|---|
| *PXN* | Paxillin | 5.4 |
| *ZEB1* | Zinc finger E-box-binding homeobox 1 | 5.1 |
| *NECAP2* | Adaptin ear-binding coat-associated protein 2 | 4.2 |
| *TPBG* | Trophoblast glycoprotein | 4.1 |
| *LMBD1* | Probable lysosomal cobalamin transporter | 3.5 |
| *MRPL12* | 39S ribosomal protein L12, mitochondrial | 2.4 |

| | | |
|---|---|---|
| *DMAP1* | DNA methyltransferase 1-associated protein 1 | 2.4 |
| *CARS2* | Probable cysteine--tRNA ligase, mitochondrial | 2.4 |
| *TGOLN2* | Trans-Golgi network integral membrane protein 2 | 2 |
| *UFSP2* | Ufm1-specific protease 2 | -3.3 |
| *MAP1S* | Microtubule-associated protein 1S | -3.8 |
| *SLC39A7* | Zinc transporter SLC39A7 | -4.1 |
| *ATPAF2* | ATP synthase mitochondrial F1 complex assembly factor 2 | -4.4 |
| *HNRNPA0* | Heterogeneous nuclear ribonucleoprotein A0 | -4.6 |
| *BUB1B* | Mitotic checkpoint serine/threonine-protein kinase | -4.7 |
| *GOLPH3L* | Golgi phosphoprotein 3-like | -5.2 |

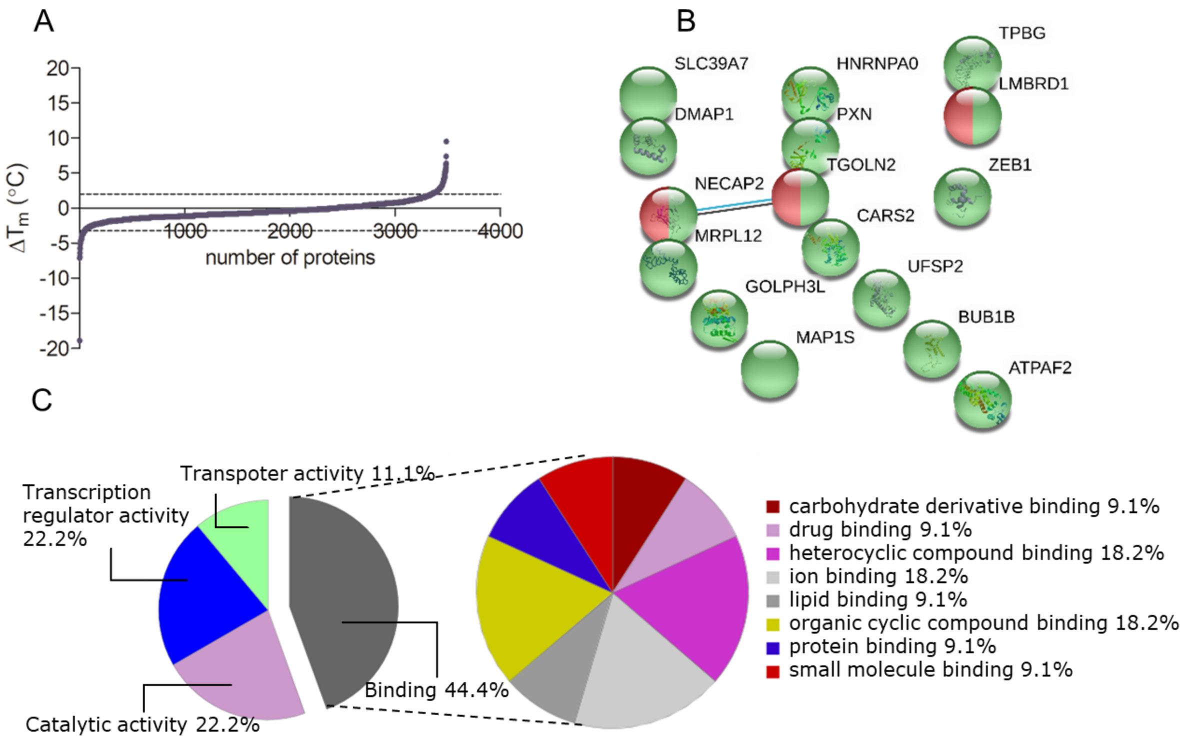

*Figure S3 A) Distribution of the melting temperature shifts for CT and two standard deviation range (dashed line); B) STRING functional enrichment analysis of the candidate targets of CT. The enriched cellular component were the intracellular organelle (GO: 0043229 in green) and the clathrin-coated vesicle (GO:0030136 in red), both with a FDR of 0.0314; C) Pie chart displaying the molecular function of the candidate targets of CT obtained by Panther classification syste. On the left, is reported the zoom of the biding activity percentage (detached from the smaller pie chart) with related legend and percentage for each function contributing to the binding GO term.*

*Table S8 Values of the single-point Km ATP assay for PI3KC2A with ATP concentration 200 μM*

| PI3KC2A | HTRF (Counts) | Mean (Counts) | Activity (% Control)* | Mean (% Control) | SD** |
|---|---|---|---|---|---|
| 25-HC @ 10μM | 1561.3 | 1351 | 100 | 103 | 4 |
| | 1141.6 | | 105 | | |
| Control (Plus Enzyme) | 1652.1 | 1548 | 99 | 100 | 3 |
| | 1783.8 | | 97 | | |
| | 1456.7 | | 101 | | |
| | 1298.8 | | 103 | | |
| Control (No Enzyme) | 9317.4 | 9385 | 1 | 0 | 1 |
| | 9452.6 | | -1 | | |

* % Control = ((sample - mean no enzyme)/(mean plus enzyme - mean no enzyme))*100
** NB. Where n = 2, the value reported here is actually range / √ 2

*Table S9 Values of the single-point Km ATP assay for PIPK1A with ATP concentration 200 μM*

| PIP5K1A | HTRF (Counts) | Mean (Counts) | Activity (% Control)* | Mean (% Control) | SD** |
|---|---|---|---|---|---|
| 7-KC @ 10μM | 1323.3 | 1342 | 104 | 104 | 0 |
| | 1360.8 | | 103 | | |
| Control (Plus Enzyme) | 1341.7 | 1596 | 104 | 100 | 3 |
| | 1538.5 | | 101 | | |
| | 1590.5 | | 100 | | |
| | 1911.5 | | 95 | | |
| Control (No Enzyme) | 8767.3 | 8469 | -4 | 0 | 6 |
| | 8170.5 | | 4 | | |

* % Control = ((sample - mean no enzyme)/(mean plus enzyme - mean no enzyme))*100
** NB. Where n = 2, the value reported here is actually range / √ 2

*Table S10 Values of the single-point Km ATP assay for PHKG2 with ATP concentration 200 μM*

| PHKG2 | Counts | Mean (Counts - Blanks) | Activity (% Control) | Mean | SD* |
|---|---|---|---|---|---|
| 7-KC @ 10μM | 5065 | 4148 | 93 | 86 | 11 |
| | 4325 | | 78 | | |
| CONTROL | 5852 | 4869 | 109 | 100 | 7 |
| | 5373 | | 99 | | |
| | 5094 | | 93 | | |
| | 5345 | | 99 | | |
| BLANK | 139 | / | / | / | / |
| | 956 | | / | | |

* NB. Where n = 2, the value reported here is actually range / √ 2

*Table S11 Values of the single-point Km ATP assay for CDK9 with ATP concentration 45 µM*

| CDK9 | Counts | Mean (Counts - Blanks) | Activity (% Control) | Mean | SD* |
|---|---|---|---|---|---|
| 4β-HC @ 10µM | 7544 | 6615 | 123 | 122 | 2 |
| | 7409 | | 121 | | |
| CONTROL | 6660 | 5420 | 107 | 100 | 5 |
| | 6242 | | 99 | | |
| | 5965 | | 94 | | |
| | 6261 | | 100 | | |
| BLANK | 1311 | / | / | / | / |
| | 413 | | / | | |

* NB. Where n = 2, the value reported here is actually range / √ 2

#### Materials and Methods

##### 4β - Hydroxycholesterol synthesis

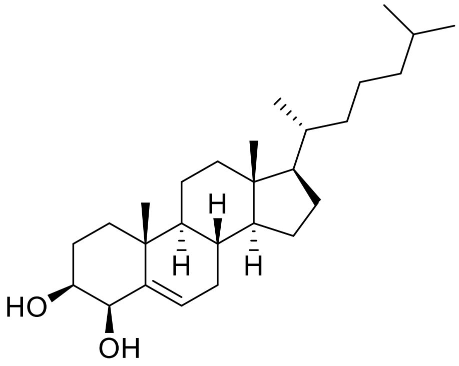

4β -Hydroxycholesterol (3β,4β-Dihydroxy-5-cholestene) was synthesized as previously described by Ma and Choi[2]. Experimental data was consistent with that reported in the literature.

$^{1}$H NMR (400 MHz, $CDCl_3$) δ 5.68 (dd, *J* = 4.8, 1.8 Hz, 1H), 4.13 (d, *J* = 3.4 Hz, 1H), 3.56 (dt, *J* = 11.8, 4.1 Hz, 1H), 2.12 – 1.98 (m, 2H), 1.94 – 1.80 (m, 5H), 1.67 – 1.23 (m, 12H), 1.18 (s, 3H), 1.14 – 0.96 (m, 8H), 0.91 (d, *J* = 6.5 Hz, 3H), 0.86 (dd, *J* = 6.6, 1.8 Hz, 7H), 0.68 (s, 3H).

$^{13}$C NMR (101 MHz, $CDCl_3$) δ 142.77, 128.82, 77.22, 72.50, 56.92, 56.10, 50.20, 42.32, 39.69, 39.52, 36.92, 36.19, 36.00, 35.78, 32.09, 31.83, 28.22, 28.02, 25.42, 24.26, 23.83, 22.83, 22.57, 21.06, 20.54, 18.71, 11.87.

#### $^{1}H$ NMR - 4ß-Hydroxycholesterol

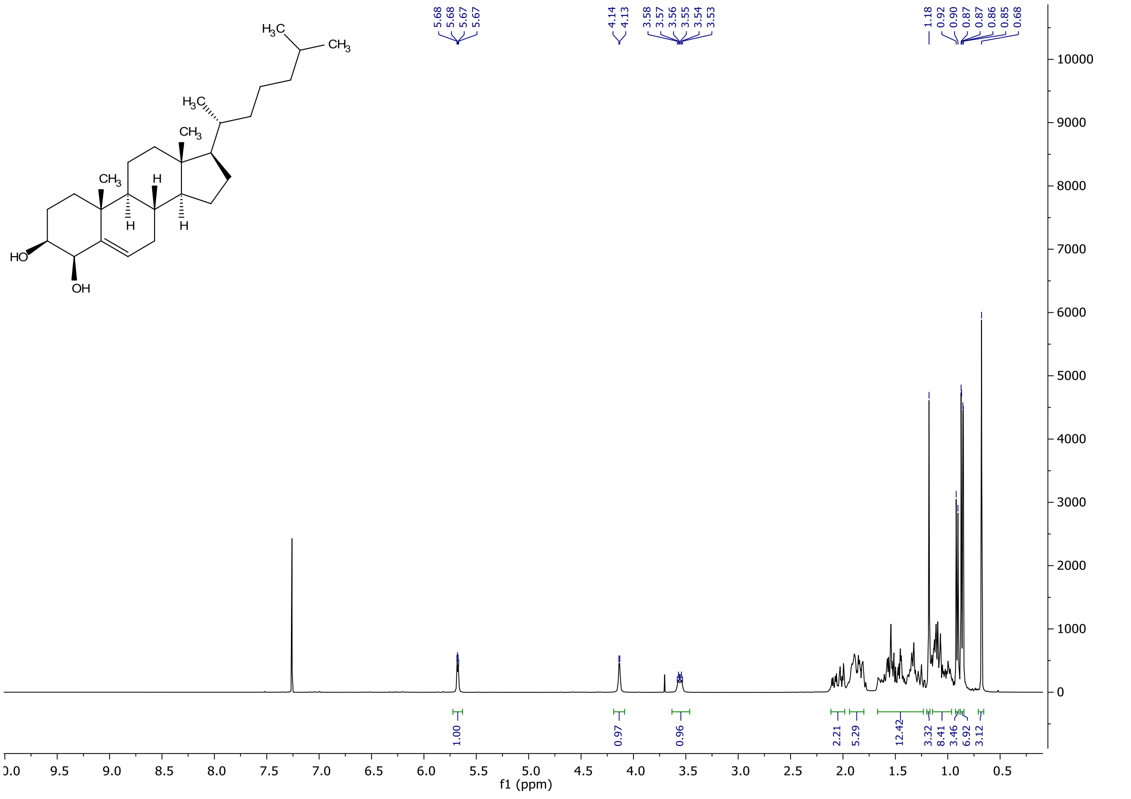

#### $^{13}C$ NMR - 4ß-Hydroxycholesterol

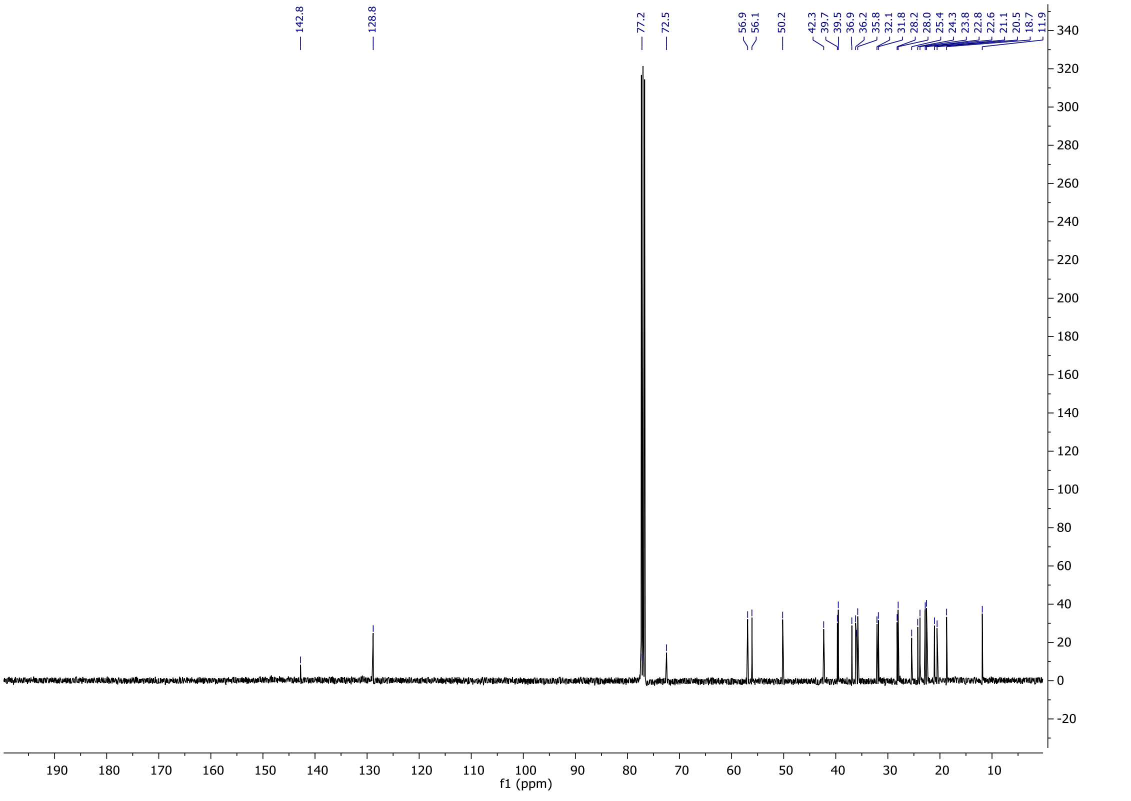

#### Cholestane-3β,5α,6β-triol synthesis

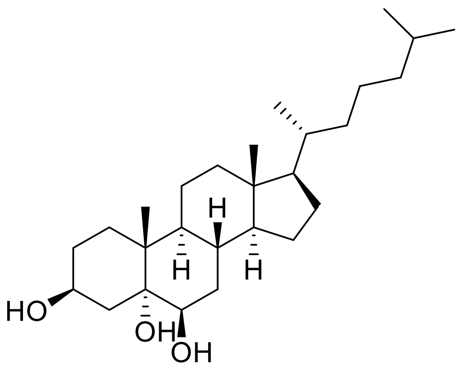

cholestane-3β,5α,6β-triol was synthesized as previously described by Li and Li[3]. Experimental data was consistent with those reported in the literature.

$^1$H NMR (400 MHz, $CDCl_3$) δ 4.10 (tt, *J* = 10.7, 5.2 Hz, 1H), 3.54 (t, *J* = 2.9 Hz, 1H), 2.09 (dd, *J* = 12.8, 11.1 Hz, 1H), 2.00 (dt, *J* = 12.6, 3.3 Hz, 1H), 1.88 – 1.66 (m, 3H), 1.64 – 1.25 (m, 20H), 1.18 (s, 3H), 1.16 – 1.06 (m, 6H), 0.90 (d, *J* = 6.5 Hz, 3H), 0.86 (dd, *J* = 6.6, 1.8 Hz, 6H), 0.68 (s, 3H).

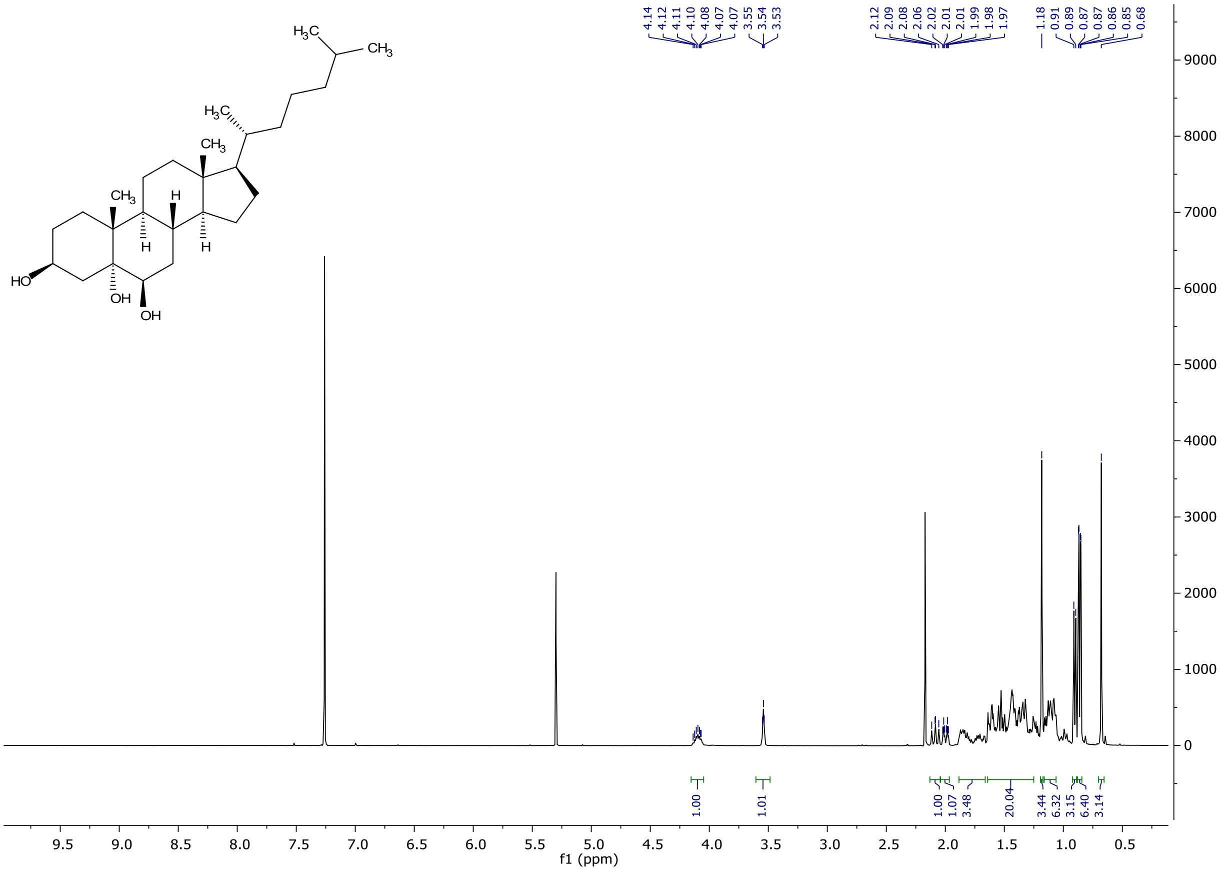

#### Reagents:

25-hydroxycholesterol was purchased from Sigma-Aldrich (Sigma-Aldrich H1015), 7-ketocholesterol from Avanti® Polar Lipids, Inc. (Avanti 700015P). All the oxysterols were dissolved in dimethyl sulfoxide (DMSO) (Sigma-Aldrich D8418) to obtain 10 mM stock solutions.

#### Cell culture

HeLa cells (ATCC Cell Biology Collection, HeLa ATCC® CCL-2™) were cultured in high-glucose Dulbecco's Modified Eagle's Medium (DMEM) (Sigma-Aldrich, D6429) supplemented with 10% fetal bovine serum (FBS) (Gibco™10270106) and 1% penicillin/streptomycin (Gibco™ 15140122). Cells were split at a ratio of 1:10 upon reaching roughly 80-90% confluence after detachment with trypsin (Gibco™ 25300054).

#### Cell lysis

Confluent cells were detached using trypsin, pooled and kept on ice. Cells were subsequently centrifuged at 4 °C for 3 min (350 × g) and cell pellets were washed 3×with ice-cold PBS (Sigma-Aldrich P4417). Cells were resuspended in ice-cold PBS containing 0.4% (v/v) Nonidet P40 Substitute (NP40) (Sigma-Aldrich 11754599001) and successively flash frozen in liquid nitrogen. Lysis of the cells was finalized by repeating four freeze-thaw cycles using liquid nitrogen, a ThermoMixer set at 23 °C (Eppendorf®) and an ice bath. The obtained lysate solution was cleared by ultracentrifugation (100000 × g) for 20 min at 4 °C. Successively, supernatants were combined and protein concentration was assessed using Bradford assay. HeLa cell extracts were diluted and further used at a final concentration of 2 mg/mL.

#### Thermal Proteome Profiling (TPP) experiments

The protocol for TPP experiments was adjusted from Reckzeh *et al*[4]. Cell extracts were thawed on ice, mixed and divided in two separate aliquots to be incubated at room temperature for 10 min with 0.1% DMSO or 10 µM of the tested oxysterol (4ß-HC; 25-HC; 7-KC or 5,6-CT). Each aliquot was split in 10 PCR-tubes which were then heated for 3 min at 10 different temperatures (37.2; 40.2; 44.9; 48.7; 51.0; 54.0; 56.9; 60.8; 64.5; 67 °C) using PCR gradient-cyclers (Eppendorf® Mastercycler®). The precipitated proteins were separated by ultracentrifugation at 100000 x g for 25 min and kept at 4 °C.

#### Protein digestion, Tandem Mass Tag (TMT) derivatization and high-pH fractionation

Supernatants from the TPP experiments were collected and diluted 1:2 with triethylammonium bicarbonate buffer (TEAB) 100 mM (Sigma-Aldrich T7408) before reduction with tris(2-carboxyethyl)phosphine hydrochloride (TCEP) 200 mM (Sigma-Aldrich C4706) for 1 h at 55 °C. Cysteine residues were successively alkylated with Iodoacetamide (IAA) (Sigma-Aldrich I1149) 375 mM for 30 min at room temperature in the dark before the addition of 6 volumes of cold acetone (Sigma-Aldrich 650501) for overnight protein precipitation at -20 °C. The following day the protein pellets were collected by centrifugation at 8000 × g at 4 °C for 10 min and resuspended in TEAB 100 mM before digestion. Lys C (WAKO 121-05063) was initially added to the samples for 4 h at 37 °C before the overnight digestion with trypsin (Sigma-Aldrich T6567) at 37 °C, with an 1:80 enzyme: substrate ratio.

After the digestion, the labelling with the TMT reagents (Thermo Fisher Scientific A34808) was performed according to manufacturer's protocol. After quenching with 5% hydroxylamine (Thermo Fisher Scientific 90115), all the 10 samples related to the DMSO incubation were pooled, as the ones incubated with the oxysterol. The resulting two aliquots were desalted by solid-phase extraction using Sep-Pak® Plus C18 cartridges (Waters WAT020515), eluting with increasing concentration of acetonitrile (40-80%) (Pierce™ 51101) with 0.1% of trifluoroacetic acid (TFA) (Honeywell™ Fluka™ 14264) and dried in SpeedVac

(Eppendorf EP022822993) at room temperature. Samples were checked for peptide concentration before injecting 30 µg in the UHPLC system for the high-pH fractionation (Dionex Ultimate 3000 system U3000, combined with an automated autosampler/fractionator). The separation of the peptides was carried out at a costant flowrate of 2.5 µl $min^{-1}$ on a CSH C18 Acquity UPLC M-Class Peptide column, 130 Å, 1.7 µm, 300 µm x 150 mm (Waters 186007563) using a 100 min linear gradient from 5 to 35% of mobile phase B (acetonitrile) with a subsequent 15 min gradient to 70%, before 5 min re-equilibration with 95% of mobile phase A (5mM ammonium bicarbonate (Supelco 5.33005), pH 10). Sixty time-based fractions were collected and supplemented with iRT peptides (Biognosys Ki-3002) in 2% acetonitrile with 1% TFA, for further check on the chromatographic performance of the liquid chromatography- mass spectrometry (LC-MS) system. The collected fractions were pooled in twenty-eight fractions before clean-up by EvoTip (EvoSep EV2003) according manufacturer's instructions.

#### LC-MS analysis

For the MS analysis, the EvoTips were loaded onto an Evosep One module (Evosep EV-1000) coupled to a Q-Exactive HF-X mass spectrometer (ThermoFisher Scientific). The loading of the peptides onto the EASY-Spray™ C18 column, 2 µm, 100 Å, 75 µm x 15 cm (ThermoFisher Scientific ES804) was performed using the "30 samples per day" standard method designed by Evosep. Peptides eluted over a 44 min gradient ranging from 5% to 90% acetonitrile with 0.1% formic acid (Pierce™85174). The MS acquisition was performed in data dependent-$MS^2$ mode, with fragmentation of the 20 most abundant ions detected in the full MS scan (Top20 method). Full MS spectra were acquired in the 350-1500 *m/z* range, with a resolution of 120000 and an automatic gain control (AGC) *target* of $3 \times 10^6$ or maximum ion injection time (IT) of 50 ms. The $MS^2$ spectra were acquired in the 350-1500 *m/z* range with the fixed first mass set to 110 *m/z* to ensure the acquisition of the TMT reporter ions and isolation window of 1.2 *m/z*. The resolution was set to 45000, AGC target to $1 \times 10^5$ or maximum IT 96 ms, normalized collision energy to 32 and dynamic exclusion set to 20 s. Minimum signal intensity was set to $1 \times 10^5$ and singly charged ions and ions with a charge state greater than 6 or unknown were excluded. The MS performances were constantly monitored by the analysis of an in-house standard of HEK293-cell lysate as quality control at the beginning of each sample set.

#### Cellular Thermal Shift Assay (CETSA) experiments and isothermal dose-response fingerprinting (ITDRF) experimets

For the CETSA experiments HeLa cell lysates were thawed on ice, separated in the two aliquots to be incubated for 10 min at room temperature with either 0.1% DMSO or with 10 µM of the tested oxysterol, and heated at 9 different temperature (37.2; 40.2; 44.9; 48.7; 51.0; 54.0; 56.9; 60.8; 64.5 °C) for 3 min using PCR gradient-cyclers.

In the ITDRF experiments HeLa lysates were thawed on ice and separated in 9 aliquots to be incubated for 10 min at room temperature with either 0.1% DMSO or with decreasing concentration of 4β-hydroxycholesterol (20; 10; 5; 2.5; 1.25; 0.6; 0.3 µM). All the aliquots were successively heated for 3 min at 51 °C using a PCR cycler.

For both CETSA and ITDRF, the removal of the precipitated proteins from the solution was performed by ultracentrifugation at 4 °C with a speed of 30000 rpm for 25 min followed. The cleared supernatants were collected and further analyzed by Western Blot.

#### Sodium dodecyl sulfate-polyacrylamide gel electrophoresis (SDS-PAGE) and Western Blot analysis

Lysate from the CETSA and ITDRF experiments were mixed with the loading buffer solution (100 mM Tris–HCl (pH 6.8) (Sigma-Aldrich T5941), 4% (w/v) SDS (Sigma-Aldrich 1.12533), 0.1% (w/v) bromophenol blue (Sigma-Aldrich B5525), 20% (v/v) glycerol (Sigma-Aldrich G5516), 200 mM DL-Dithiothreitol (Sigma Aldrich 43819) in 4:1 ratio (v/v)), heated at 95°C for 5 min for complete protein denaturation. Samples were then loaded onto 4–20% TGX Stain-Free™ Protein Gels (Bio-Rad 4568093) and run at 120 V for 1 hour. Protein transfer from gels to polyvinylidene difluoride (PVDF) membranes was carried out using the Trans-Blot Turbo Transfer Kit (Bio-Rad 1704274) according to the manufacturer's instructions. Membranes were subsequently incubated with the blocking solution (5% Skim Milk Powder (Fisher Scientific 10651135) added to tris-buffered saline (TBS) solution (150 mM NaCl (Sigma-Aldrich S7653), 50 mM Tris pH 7.6, 0.1% Triton™ X-100 (Sigma-Aldrich T8787)) before the incubation with the primary antibody overnight at 4 °C. For VPS51 the used primary antibody was Anti- VPS51 Rabbit anti-Human, Polyclonal (Invitrogen™ PA-58869) and for NECAP2 it was used Anti- NECAP2 Rabbit anti-Human, Polyclonal (Sigma-Aldrich HPA028077)

The following day membranes were incubated for 4 h with the HRP-conjugated secondary antibody Goat Anti-rabbit IgG (Invitrogen™31460), which was diluted 1:10000 in blocking solution. Successively, the SuperSignal™West Femto Chemiluminescent substrate (Thermo Scientific™ 10391544) was added to the membranes before imaging with a ChemiDoc XRS+ System (Bio-Rad). Image acquisition and quantification of the chemiluminescent intensities of the bands were performed by Image Lab™ version 6.0.1 (Bio-Rad).

#### Data Analysis

Peptide and protein identification and quantification, as well as the generation of the melting curves from the TPP experiments were carried out according the protocol described in Reckzeh et al.[4].

Briefly, MaxQuant was used to process the raw data from the MS runs, setting all the TMT10 labels as reporter ion MS2 for lysine and N-terminus with mass tolerance 0.01 Da. Methionine oxidation and acetylation of the protein N-terminal were set as variable modifications, while carbamidomethylation was the fixed modification. Maximum number of modifications per peptide was set at 5, while the minimum peptide length was set at 7, with 7 as maximum charge. Specific digestion with LysC and Trypsin was allowed for maximum 2 missed cleavages. Peptide-spectrum match (PSM) and protein false discovery rates (FDR) were 0.01 with at least 1 minimum razor + unique peptides. Peptide sequences were searched against UniProtKB/ TrEMBL human proteome[5] fasta file (https://www.uniprot.org/proteomes/ accessed on the 08/08/2018).

The protein group files generated from the MaxQuant analysis were processed in the Excel Macro "TPP_Makro_1.0" (freely available on http:// www.mpi-dortmund.mpg.de/forschung/chemische-biologie/janning), where the reporter ion intensities at the different temperatures for each protein are fit according the equation:

$$y = bottom\,plateau + \frac{(top\,plateau - bottom\,plateau)}{1 + e^{-\left(\frac{a}{Temp}\right)-b}}$$

Top plateau is set to one, due to the normalization of all intensities to the lowest temperature, while the variable Temp is the temperature, the bottom plateau is a protein-specific constant regarding the maximal denaturation and a and b are constants which describe the curve progression.

Melting temperatures were calculated from the first derivative of the curves and proteins were classified according the criteria described in Reckzeh et al.[4]

The results generated from the macro have been further filtered according the following procedure:

All the curves without thermal shift in all the three replicates and protein with thermal shift greater than 30 ˚C were excluded (moved in the No sheet of the macro). The median of the thermal shifts, together with the standard deviation are calculated for the remaining proteins and plotted in the graphs vs the protein number (Stat&Graphs sheet). Thermal shift significance was set as greater than 95.5% distribution probability, therefore shifts which were greater than the upper limit (set as median + 2 times the standard deviation) and smaller than the lower limit (set as median – 2 times the standard deviation) were considered significant, in line with the approach previously reported from Justice et al.[6]

Proteins with significant thermal shift were further validated according the bottom plateau of the curves and the direction of the thermal shifts in all the three replicates. Proteins with bottom plateau below 0.5 and comparable thermal shift in all the replicates, were assigned as putative targets and moved in the Yes sheet of the macro, while protein bottom plateau greater than 0.5 or with thermal shifts in the same direction for two out of the three replicates were assigned in the Maybe sheet of the macro. In the No sheet were moved proteins with different thermal shift direction in all the replicates and the protein identified as contaminant or reverse sequence by MaxQuant.

#### Data analysis and statistics

Western blot data generated from Image Lab™ reports, were further analyzed using Microsoft Excel for normalization (version 16) and Prism 5 (Graphpad) for plotting melting curves.

Functional classification and enrichment analysis of putative protein targets was performed with the Enrichr, an enrichment analysis web server[7][8], PANTHER tools[9] filtered as protein molecular functions (http://pantherdb.org accessed on the 11/09/2020) and STRING[10] set with medium confidence interaction score (https://string-db.org accessed on the 28/09/2020). Venn diagram of the hits found for each oxysterol was created with VENNY 2.1 (https://bioinfogp.cnb.csic.es/tools/venny/index.html accessed on the 11/09/2020)[11]

#### Kinase profiling

The validation of selected kinases found as putative targets (CDK9, PHKG2, PIK3C2A and PIP5K1A) was performed by Eurofins with single-point [Km ATP] assays (Eurofins Discovery). The assays for CDK9 and PHKG2 were radiometric assays, while the assays for PIK3C2A and PIP5K1A were homogeneous time resolved fluorescence (HTRF) assay.

The radiometric assays provided the incubation of the selected kinase with 8 mM MOPS pH 7.0, 0.2 mM EDTA, 100 µM KTFCGTPEYLAPEVRREPRILSEEEQEMFRDFDYIADWC for CDK9 or 250 µM KKLNRTLSFAEPG for PHKG2, 10 mM MgAcetate and [gamma-33P]-ATP (Km specific activity and concentration). The reaction is initiated by the addition of the Mg/ATP mix. After incubation for 40 minutes at room temperature, the reaction is stopped by the addition of phosphoric acid to a concentration of 0.5%. 10 µL of the reaction is then spotted onto a P30 filtermat and washed four times for 4 minutes in 0.425% phosphoric acid and once in methanol prior to drying and scintillation counting.

The HTRF assays provided the incubation of the selected kinase in assay buffer containing 25 µM phosphatidylinositol (for PI3KC2A) or phosphatidylinositol 4-phosphate (for PIP5K1A)/75 µM phosphatidylserine and Mg/ATP (Km concentration). The reaction was initiated by the addition of the ATP solution. After incubation for 30 minutes at room temperature, the reaction was stopped by the addition of stop solution containing EDTA and biotinylated phosphatidylinositol-3 phosphate (for PI3KC2A) or biotinylated phosphatidylinositol 4,5-bisphosphate (for PIP5K1A). Finally, detection buffer was added, which contains europium-labelled anti-GST monoclonal antibody, a GST-tagged PX (for PI3KC2A) or pH (for PIP5K1A) domain and streptavidin allophycocyanin. The plate is then read in timeresolved fluorescence mode and the HTRF signal is determined according to the formula HTRF = 10000 x (Em665nm/Em620nm).

[11] J. C. Oliveros, "VENNY. An interactive tool for comparing lists with Venn Diagrams," **n.d.**
